# The synergistic effects of plants and nitrogen on microbial hitchhiking

**DOI:** 10.1101/2022.05.09.491057

**Authors:** Zhibin Liu, Ziyuan Wang, Qini Xia, Qin Zhou, Xiaobo Wu, Wenqing Kong, Wenyan Lei, Jiayi Zeng, Chao Liu, Yongfeng Wang, Wei Chang, Zhi Li, Yi Yang, Liang Yang, Xiao Tan

**Affiliations:** Key Laboratory of Bio-Resource and Eco-Environment of Ministry of Education, College of Life Sciences, Sichuan University, Chengdu 610064, Sichuan, PR China; State Key Laboratory of Hydraulics and Mountain River Engineering, Sichuan University, Chengdu 610065, Sichuan, PR China; College of Water Resource & Hydropower, Sichuan University, Chengdu 610065, Sichuan, PR China; Institute of Environment and Ecology, Institute of Environmental Health and Ecological Security, Jiangsu University, 301 Xuefu Road, Zhenjiang, 212013, PR China.; Guangdong Provincial Key Laboratory of Silviculture, Protection and Utilization, Guangdong A; Vegetable Germplasm Innovation and Variety Improvement Key Laboratory of Sichuan Province/Horticulture Research Institute, Sichuan Academy of Agricultural Sciences, Chengdu 610066, PR China

**Keywords:** microbial hitchhiking, rhizosphere microbes, nitrogen fertilizer

## Abstract

Microbial hitchhiking demonstrates that some nonmotile microbes utilize trans-species motility to traverse their environment; however, whether driving forces, such as plants and nitrogen, affect microbial hitchhiking is not clear. In our study, we explored the effects of plants and nitrogen fertilizer on *Bacillus*-hitchhiking by setting filter membranes and different nitrogen fertilizer concentration gradients. In the experimental treatment, we added a filter membrane to the soil to prevent hitchhiking. In the absence of plants, nitrogen alone had little influence on motile bacteria and hitchhiking. However, *Bacillus* contents were significantly impacted by the nitrogen concentration when the plants were rooted, leading to a great variation in cell motility function according to the functional analysis in the soil microbial community. After applying the filter membrane, there were no significant differences in *Bacillus* contents, microbial community structure or cell motility functional abundance, which illustrated that hitchhiking impacted the microbial community. Our analysis of co-occurrence between bulk soil motile bacteria (*Bacillus*) and rhizosphere bacteria also confirmed this. The correlation between bulk soil motile bacteria and the rhizosphere microbial community was strong in the groups with suitable nitrogen concentrations without filter membranes and was weak at all nitrogen levels in the no-membrane treatments. Thus, we concluded that plants and different nitrogen doses synergistically altered the soil microbiome by hitchhiking, whose effect depends on nitrogen.

## 1. Introduction

Cell motility is responsible for diverse and interesting survival behaviors(Kearns, 2010; Lambert et al., 2019). Many free-living motile microbes use motility machinery to navigate their surroundings in search of optimum conditions(Lambert et al., 2019). Motile microbes, which can be attracted or repelled by specific stimuli on surfaces, can travel by creeping, gliding, rolling, swarming, or twitching, among other motile behaviors. In addition to the microbe motility machinery, a novel mode of dispersal used by many microbes, hitchhiking on motile microbes, was recently reported (Jarrell and McBride, 2008; Kearns, 2010; Schäfer et al., 1998). Hitchhiking enables nonmotile microbes to navigate their world by successfully using the motility machinery of their motile partner (Muok et al., 2021). Transport by hitchhiking occurring among bacteria is found in soils, plant tissues, abiotic surfaces, and human tissues(Muok et al., 2021; Muok and Briegel, 2021). Various species of bacteria comigrate across air gaps, which allows nonmotile bacteria to access novel microhabitats(Warmink and van Elsas, 2009).

A previous study showed that plants shape rhizosphere microbiome structures by hitchhiking through motility of *Bacillus* subtilis, which thrives near plant roots and moves toward the root(Muok et al., 2021; Muok and Briegel, 2021). Bacterial motility has two modes of flagellar-mediated motility, swimming and s warming, and can also move through sliding(Muok et al., 2021). It was reported that *Bacillus* was able to transport Streptomyces spores to plant tissues. However, previous studies on how *Bacillus* hitchhiking affected the soil microbiome were carried out in an ideal laboratory environment without simulating the real soil ecological environment. At the same time, in addition to Streptomyces, it was not clear whether the other bacteria could be carried by *Bacillus*.

Some physical and chemical properties of soil, including pH and water content, determine the composition of the microbial community (Garrett et al., 2008; Lambert et al., 2019; Tsagkari and Sloan, 2018). Nitrogen, by impacting physical and chemical properties, acts directly as a key factor influencing microbial community structure(Good and Beatty, 2011; Kavamura et al., 2018; Sun et al., 2020; Yuan et al., 2015; Zhou et al., 2017). A recent study showed that various kinds of bacteria were involved in community interactions via a community co-occurrence network analysis under different nitrogen treatments, but no sources of the microbes responsible for the changes in the microbial community were identified (Hartman and Tringe, 2019). Moreover, it did not reveal whether hitchhiking played an important role in the changes in the microbial community.

To simulate the natural soil environment and explore the influence of nitrogen on microbial hitchhiking, we planted *spinach* under wild conditions by applying different nitrogen concentrations and set up a blank control group (no plants). In addition, more kinds of bacteria can be identified to determine whether they are related to hitchhiking. To explore the impact of hitchhiking on the community, we set the filter group to isolate bacterial movement. Then, high-throughput sequencing of the obtained amplicons was performed to illustrate the changes in the bulk soil and rhizosphere microbial communities under different nitrogen concentrations and with or without *spinach* planting. Furthermore, the mechanism responsible for the effects of filter membrane applications near rhizosphere soil exposed to various nitrogen concentrations and with or without *spinach* planting on plant growth was investigated. We assessed the current cooccurrence network and studied the activation of the symbiotic mode under various nitrogen concentrations and with or without *spinach* cultivation, and we found that nitrogen concentration and with or without plants influenced both motile and their associated nonmotile bacteria together.

## 2 Materials and methods

### 2.1 Sites and experimental setup

To investigate the coupled effect of fertilization and microorganism on plant growth, a pot experiment was conducted in Shuangliu District, Chengdu, China (102°52’11’’E, 30°33’21’’N). The Shuangliu District belongs to subtropical humid monsoon climate zone with an annual average air temperature range of 15.6L to 16.9L, an annual average precipitation range of 759.1 mm to 1,155.0 mm, and an annual average sunshine duration range of 825.0 h to 1,202.9 h. In this region, the soil is yellow loam and mostly slightly acidic.

The pot experiment was conducted from October 2019 to February 2020. Three nitrogen levels (no addition of extra nitrogen, addition of 20% extra nitrogen, addition of 40% extra nitrogen), two planting conditions (with/without plants), and two coating conditions (with/without filter membrane) were considered, resulted in 12 treatments. Each treatment has 3 replications. A plot (4 × 7.4 m in size) was selected from a greenhouse in the experimental site, consisting of three subplots (3 × 1.8 m in size). Each subplot has 12 sampling points, distributed in 3 rows and 4 columns (each column represents one treatment). Intervals with a width of 30 cm between each sampling point and buffer zones with a width of 30 cm between the side row/column and the plot boundary were set to reduce the disturbance (Supplementary Information). The soil used in this experiment was taken from each sampling point (diameter: 20 cm, depth: 20 cm), then air-dried, ground to pass through a sieve of 2 mm, and mixed well (guarantee the same soil texture in all treatments). Urea, nitro-compound fertilizer, and potassium sulfate were applied as base fertilizers at rates of 226.5 kg/ha, 400.6 kg/ha, and 120 kg/ha, respectively, in accordance with local agricultural practices. Afterwards, urea was added to achieve different nitrogen concentrations: base fertilizer (H1), base fertilizer + 70.5 kg/ha urea (H2), base fertilizer + 141 kg/ha urea (H3). The well-mixed soil of each nitrogen concentration was then packed into 12 fiber pots (diameter: 20 cm, height: 20 cm, 8 kg soil for each pot, growth of spinach roots would be not limited in the pot in such size) which allows mass and energy exchange between the soil inside and outside of the pot. Taking H1 as an example, half of the 12 pots (6 pots) were coated by 0.45 μm filter membrane (Fig. 1B). Then, all the prepared pots were carefully embedded into sampling points with no gaps between pots and surrounding soils. The soil surface in pots was flush with the ground. The spinach (Spinacia oleracea L.) seeds were soaked in distilled water for 6 h and then sown into half of the coated pots (3 pots) and half of the uncoated pots (3 pots). Each column refers to a treatment in a subplot, details are shown in Supplementary Information. Excessive seedlings were removed when the third true leaf emerged, and one seedling was left in each pot. Water was supplied in accordance with local water management practices. The same operations were repeated for the H2 and H3. All abbreviations in the text and corresponding treatment are shown in Fig. 1C.

**Figure 1.**
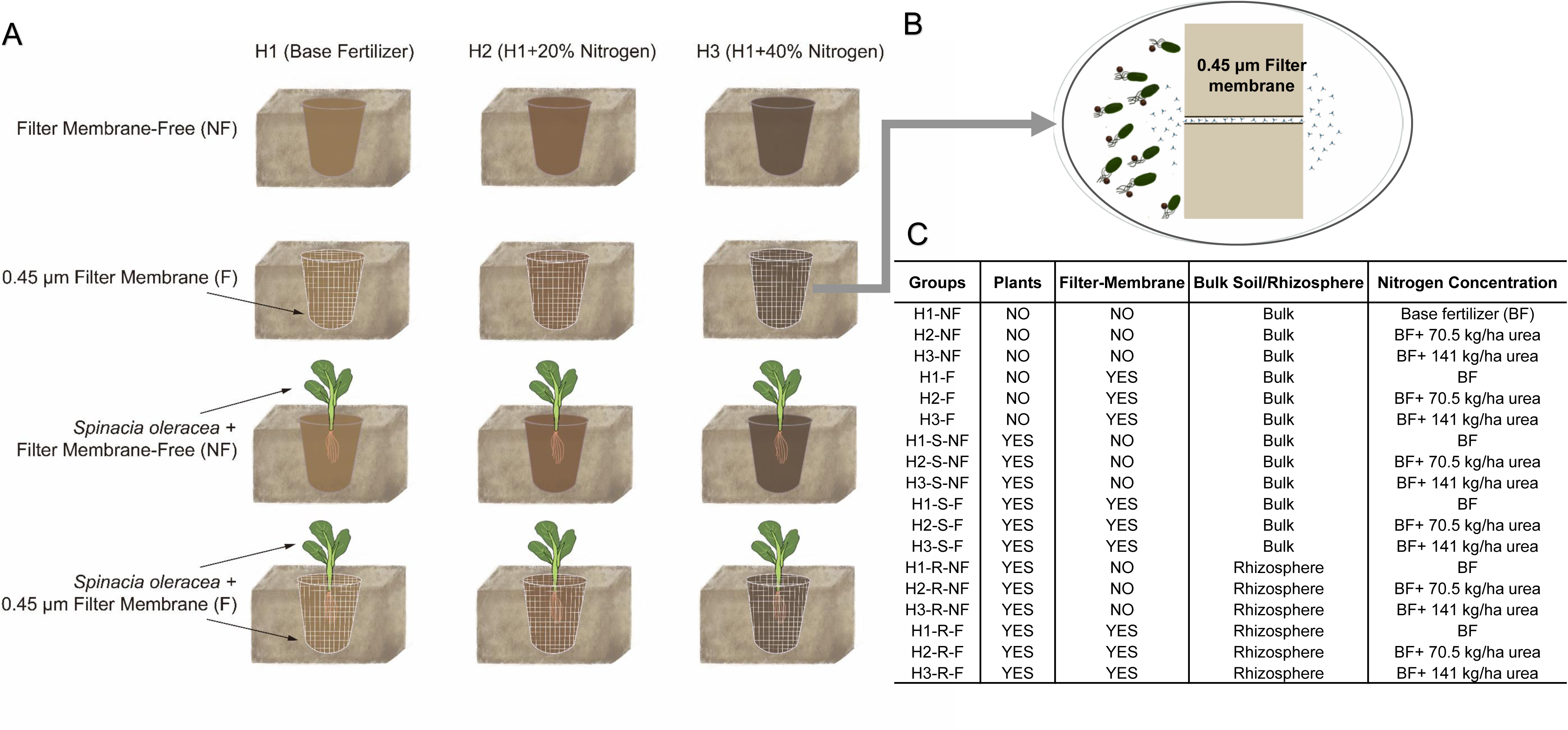
(**A**)The plots had the following soil treatment conditions with three different nitrogen fertilizer concentrations (H1–3): natural, untreated soil (NF) with plants (H1-NF, H2-NF, and H3-NF), filter membrane (F) used to separate the rhizosphere soil from the outer bulk soil (H1-F, H2-F, and H3-F). There were two treatments for each area, a no spinach and no membrane addition treatment and a spinach and (**B**) 0.45-μm filter membrane addition. (**C**) All abbreviations in the text and corresponding treatment are shown in (C).

### 2.2 DNA extraction, PCR amplification, and Illumina NovaSeq 6000 sequencing

Sample collection was conducted in February 2020. Subplots (0.5 m × 0.5 m) were randomly selected from each treatment site. Soil samples (0–15-cm depths) were drilled using a soil sampler with a diameter of 3.5 cm. Each soil sample was cut vertically in the middle, put into a sterile self-sealing bag, and transported on ice. They were then stored in a refrigerator at −80°C for microbial sequencing. Microbial DNA was extracted using a HiPure Soil DNA Kit (or HiPure Stool DNA Kit) (Magen, Guangzhou, China) in accordance with the manufacturer’s protocol. The 16S rDNA target regions of the ribosomal RNA gene were amplified by PCR using the following amplification protocol: 94°C for 2 min, followed by 30 cycles at 98°C for 10 s, 62°C for 30 s (except for 16S V4, which was 55°C for 30 s), and 68°C for 30 s, with a final extension at 68°C for 5 min. PCRs were performed in triplicate. Each 50-μL mixture contained 5 μL of 10× KOD Buffer, 5 μL of 2 mM dNTPs, 3 μL of 25 mM MgSO4, 1.5 μL of each primer (10 μM), 1 μL of KOD polymerase, and 100 ng of template DNA. The PCR reagents were purchased from TOYOBO, Japan. Amplicons were extracted from 2% agarose gels, purified using an AxyPrep DNA Gel Extraction Kit (Axygen Biosciences, Union City, CA, USA) in accordance with the manufacturer’s instructions, and quantified using an ABI StepOnePlus Real-Time PCR System (Life Technologies, Foster City, CA, USA). Purified amplicons were pooled in equimolar ratios and paired-end sequenced (PE250) on an Illumina platform using standard protocols.

### 2.3 Community composition analysis

The representative OTU sequences were classified into organisms by a naive Bayesian model using RDP classifier[15] (version 2.2) based on SILVA[16] database (version 132), with the confidence threshold value of 0.8 (Table S1, Table S2). The stacked bar plot of the community composition was visualized in R project ggplot2 package (version 2.2.1).

### 2.4 Statistical analyses

#### 2.4.1 Alpha- and beta-diversity analyses

Chao1 and Sob indices were calculated using QIIME (Caporaso et al., 2010) (version 1.9.1). An OTU rarefaction curve, as well as rank abundance curves, were plotted in R project ggplot2 package (version 2.2.1, URL: http://CRAN. R-project.org/package=ggplot2, URL: http://CRAN.R-project.org/package=ggplot2). An alpha-index comparison between groups was calculated using Welch’s t-tests in the R project Vegan package (version 2.5.3). Alpha-index comparisons among groups were conducted using Kruskal–Wallis H tests in the R project Vegan package (version 2.5.3).

The sequence alignment was performed using Muscle (Edgar, 2004) (version 3.8.31), and a phylogenetic tree was constructed using FastTree(Price et al., 2010) (version 2.1). Then, weighted and unweighted unifrac distance matrices were generated using the GuniFrac package (Lozupone and Knight, 2005) (version 1.0) in R project. Bray–Curtis distance matrices were calculated in the R project Vegan package (version 2.5.3). Multivariate statistical techniques, including a principal coordinates analysis (PCoA) and the Bray–Curtis distances, were generated in the R project Vegan package (version 2.5.3) and plotted in R project ggplot2 package(version 2.2.1.). Statistical analyses of Welch’s t-tests, Kruskal-Wallis H tests, and Anosim tests were calculated in the R project Vegan package (version 2.5.3).

#### 2.4.2 Functional predictions

The KEGG pathway analysis of the OTUs was inferred using Tax4Fun (version 1.0)(Aßhauer et al., 2015). Tax4Fun arranges all the samples, including SILVA species and annotated OTUs. An absolute abundance table, which is a program based on the SILVA species annotation information database, selects OTU reference species and then combines the OTU abundance table and species-gene nets. The output of each sample is the relative abundance of the KEGG Orthology pathway.

#### 2.4.3 Co-occurrence analysis

Graphical representations were generated using Cytoscape (Version 3.8.2). A bacterial co-occurrence analysis was computed in the Vegan package of R 4.0.2. One network per treatment combination was calculated to compare the effects of different nitrogen concentrations. The input data for the visual comparison were mean abundances over the three field replicates per time point collapsed into taxonomic families. For statistical evaluation, topological networks were computed for each plot, and Cytoscape (Version 3.8.2) was used to generate graphical representations of the resulting consensus networks. The average degree, average clustering coefficient, and average path length were computed and compared statistically using distance and Pearson’s correlation values to show the co-occurrence networks of each bacterial class.

## 3. Results

### 3.1 Bulk and rhizosphere soil compositional differences

Major microbial classes in the bulk soil without plants were *Gammaproteobacteria*, *Alphaprotebacteria*, and *Planctomycetacia* (Fig. 2A, Table S1). The content of *Planctomycetacia* increased with the nitrogen concentration, and this phenomenon was blocked by the filter membrane. In addition, more classes were influenced by nitrogen (*Planctomycetacia*, *Verrucomicrobiae*, and *Actinobacteria*). Nevertheless, the percentages of *Planctomycetacia*, *Verrucomicrobiae*, and *Actinobacteria* did not change significantly in the filter membrane treatment. Compared with treatments without plants, *Bacteroidia* accounted for a larger proportion of the community, while Planctomycetacia accounted for a smaller proportion (Fig. 2B). In the rhizosphere microbial community, *Gammaproteobacteria* abundance in H2 was greater than that in H1 and H3, irrespective of the membrane (Fig. 2C).

**Figure 2.**
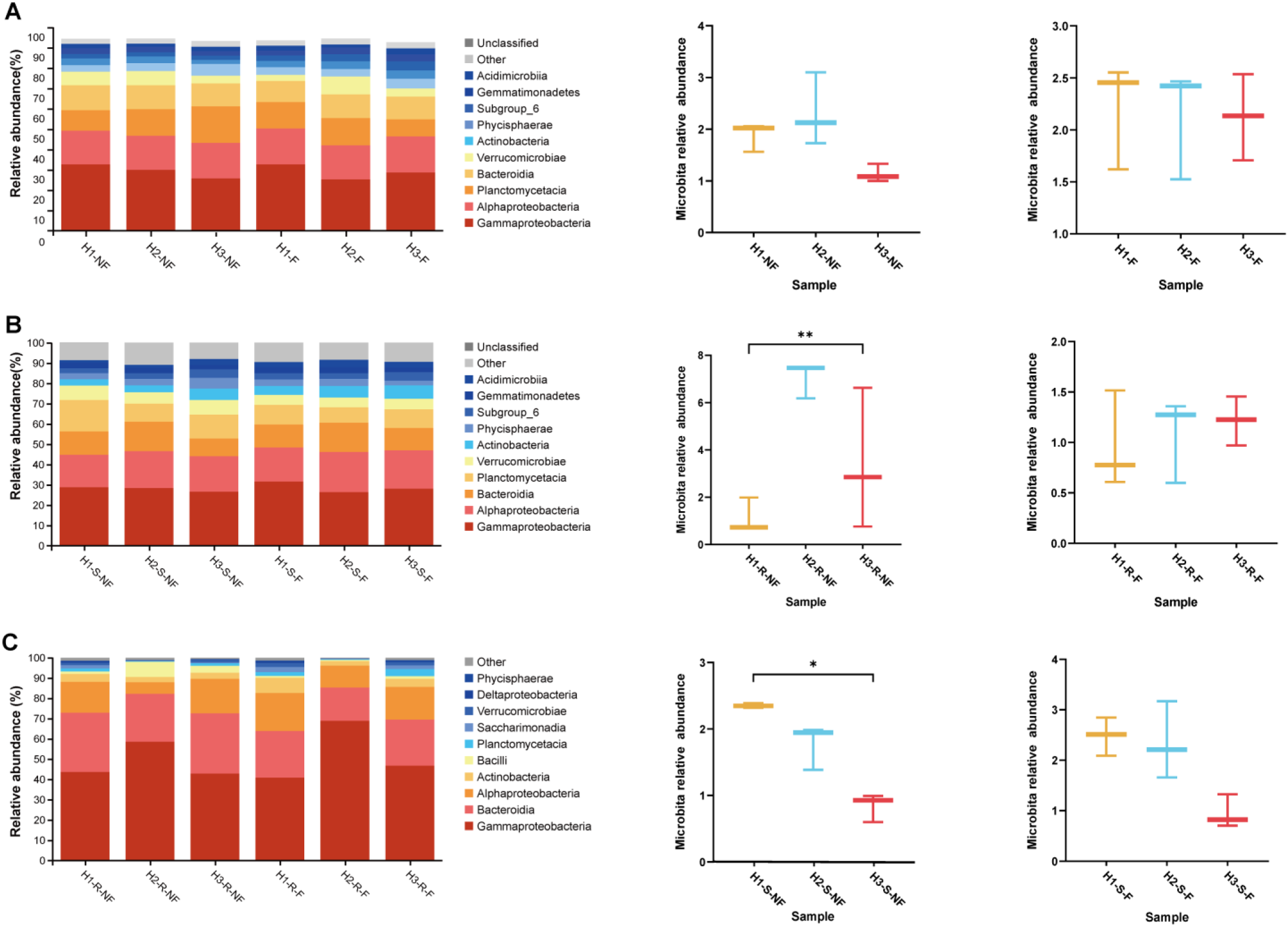
Comparison of bacterial community classification differences (class level) under different nitrogen treatments. (A, B, C) The relative abundances of Bacillus contents in soil without plants(A), without plants(B) and rhizosphere (C). H1, H2, and H3, normal, normal + 20%, and normal + 40% nitrogen fertilizer concentrations, respectively. S, bulk soil. R, spinach rhizosphere soil. F, filter membrane present, and NF, no filter membrane. * p<0.05,** p<0.01

In the soil microbial community without plants, there was no significant difference in the content of *Bacillus* regardless of the filter membrane (Fig. 2A). When there were plants in the soil, there was a significant difference in the content of *Bacillus* only if there was no filter membrane (p<0.05 without filter membrane and p=0.0714 with; Fig. 2B). In the rhizosphere microbial community, the content of *Bacillus* at different nitrogen concentrations showed a similar trend as the soil microbial community with the plants (p<0.05 without filter membranes and p=0.9286 with; Fig. 2C).

### 3.2 Microbial alpha- and beta-diversity levels

The Sob indices were used to estimate and compare the alpha-diversity of bacterial communities among different nitrogen treatments (Table 1A, Table S3). In treatments with or without the filter membrane, whether plants were added and the bulk soil/rhizosphere bacterial species diversity was not significantly different among the three different nitrogen concentrations (P> 0.05; Table 1A, Table S4A). PCoA showed a distinct clustering pattern for the microbiota in H1-NF/ H2-NF/ H3-NF(Fig. S1A), H1-S-NF/ H2-S-NF/ H3-S-NF (Fig. 3A) and H1-R-NF/H3-R-NF/H2-R-NF (Fig. 3B).

**Figure 3.**
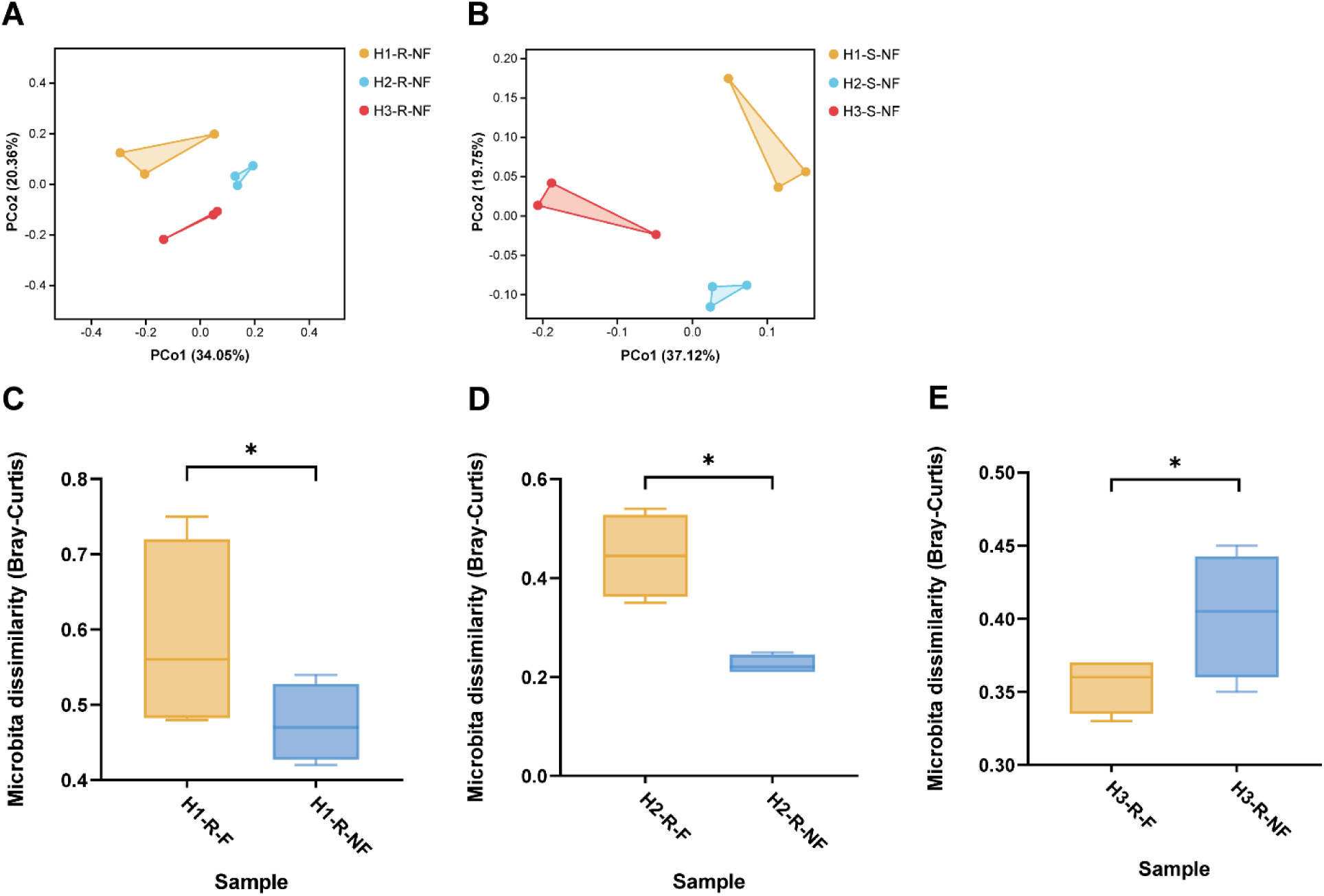
Beta-diversity levels of bulk soil and rhizosphere bacterial communities. (A) The PCoAs of soil and rhizosphere bacteria. Soil type is the main source of bacterial community changes, and each spot corresponds to different soil-type samples. (B) Significance test analysis of rhizosphere bacteria. The ends of the line represent the minimum and maximum values, and the line in the box is the middle value. H1, H2, and H3, normal, normal + 20%, and normal + 40% nitrogen fertilizer concentrations, respectively. S, spinach planted. R, spinach rhizosphere soil. F, filter membrane present, and NF, no filter membrane.

At the class level, for the bulk soil microbial community with and without plants, the bacterial beta-diversity (P = 0.003**, Table 1B; P=0.026*, Table S4B) varied greatly with nitrogen concentration. Nevertheless, there were no significant differences in beta-diversity levels among the different nitrogen concentration groups after membrane addition (P = 0.263, Table 1B; P=0.2; Table S4B). Moreover, there was no significant change in the beta diversity of the soil microbial communities between groups with and without filter-membrane (Fig. S1 B-D). A further analysis of the rhizosphere microbial colonies showed the same results as the beta-diversity analysis in the bulk soil microbial community with plants.

We discovered differences between the microbial communities at the same nitrogen concentration with and without the filter membrane treatment (Fig. 3C– E). In the bulk soil, both alpha and beta diversity showed significant differences between groups with and without filter membranes, whether there were plants or not. In a comparison of H1-R-NF and H1-R-F, although no significant difference in their alpha-diversity levels was found, there was a significant difference in their beta-diversity levels, which reflects a significant difference in their microbial community structures. When comparing H2-R-NF to H2-R-F and H3-R-NF to H3-R-F, the same result was found. Thus, although there were great effects of nitrogen on the bacterial community, the addition of the membrane counteracted the effects.

### 3.3 Abundance of function

To determine the functional changes among the different nitrogen treatments, we utilized and compared the relative functional abundances of the different groups(Table S5). In the absence of plants, nitrogen fertilizer had very little effect on cell motility abundance (Fig. S2 A). However, in H1-NF/H2-NF/H3-NF, the greatest functional abundance difference occurred in the H2 group compared with the other two treatments in both bulk and rhizosphere soils, and the functions were weakened in the membrane-addition treatments (Fig. 4A; Fig. S2 B). In particular, cell motility showed significant differences, with H2-S/R-NF showing significantly more cell motility than H1-S/R-NF and H3-S/R-NF (P < 0.05; Fig. 4 C-D). However, the level of cell motility in the membrane-addition group was not significantly different among the different nitrogen treatments. In addition, this phenomenon was not observed in soil microbial communities regardless of whether plants were grown.

**Figure 4.**
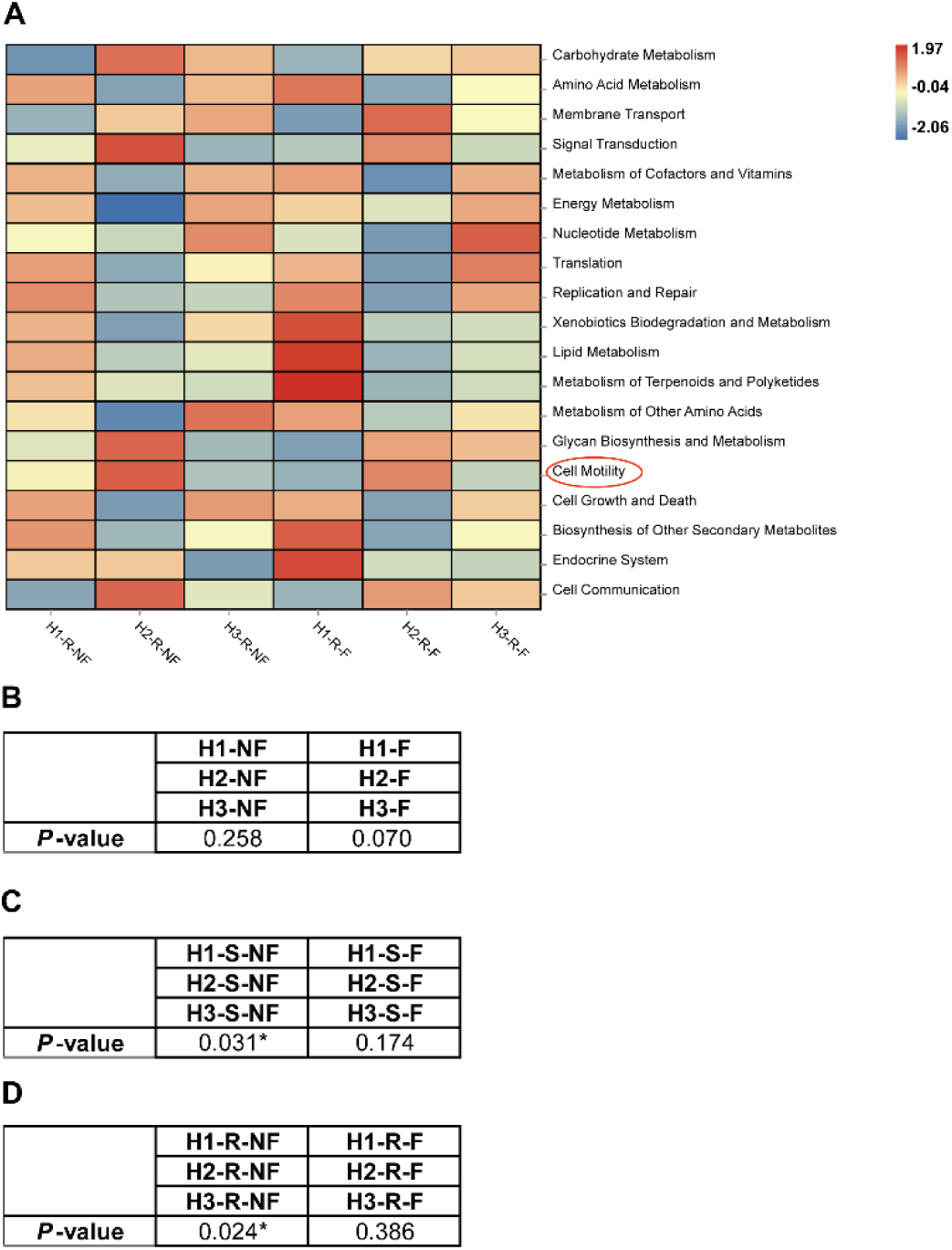
The relative abundances of the top 20 classes of the most abundant rhizosphere bacteria plotted as a functional heatmap. H1, H2, and H3, normal, normal + 20%, and normal + 40% nitrogen fertilizer concentrations, respectively(A). Cell Motility was selected and marked red, and its test in bulk soil without plants (B), with plants (C) and rhizosphere(D) and different nitrogen concentration. R, spinach rhizosphere soil. F, filter membrane present, and NF, no filter membrane.

### 3.4 Co-occurrence networks between bulk soil and rhizosphere soil microbial communities under different nitrogen concentrations

We used Cytoscape (Version 3.8.2) to draw the Co-occurrence networks and analyze the interactions of microbial communities exposed to different nitrogen concentrations. The Pearson’s and distance correlation values between classes were calculated based on their occurrences across samples using the R project Vegan package. The microorganism classes with high abundance values and high correlations between nodes were selected, and an Co-occurrence network diagram was constructed. Because certain bacteria can survive only in soil at a certain nitrogen concentration, we chose the top 19 most abundant groups for the analysis. To establish the Co-occurrence network graph, edges with Pearson’s correlation coefficients greater than 0.7 were selected.

While the number of edges in the bacteria indicating an association between bulk soil and rhizosphere microbial community increased with the nitrogen concentration, the numbers of edges under H2 and H3 were significantly greater than those under H1, which was consistent with groups having the filter-membrane treatment. In H2, without the filter-membrane treatment, the relationships between the bulk soil motile bacteria (*Bacillus*) and the rhizosphere microorganisms were significantly stronger than those in the other two groups (Fig. 5A–C). However, in groups with the filter membrane treatment, the numbers of edges indicated that associations with motile bacteria were significantly reduced in all three treatments compared with groups without filter membranes. Moreover, there were no differences among the three treatments with the filter membrane (Fig. 5D–F).

**Figure 5.**
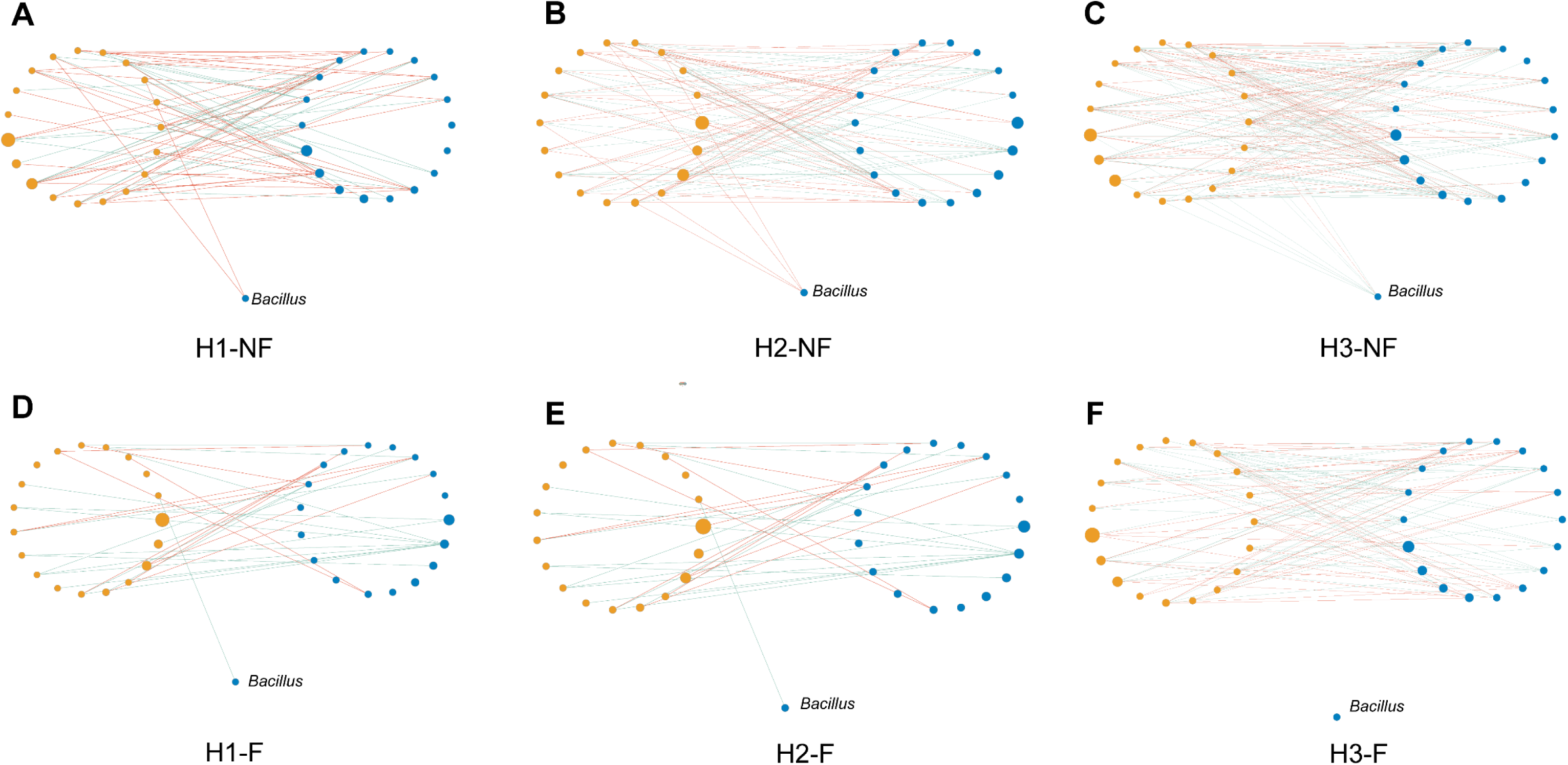
Microbial interaction networks of the different treatments. Different nitrogen treatments (A–C) without the filter membrane (NF) and (D– F) with the filter membrane (F). H1, H2, and H3, normal, normal + 20%, and normal + 40% nitrogen fertilizer concentrations, respectively. The interaction network of dominant microbiota at the class level (top 19) between bulk soil (Blue nodes) and rhizosphere (Red Nodes). The sizes of the nodes indicate the abundance levels of different bacterial classes, and the different colors indicate the corresponding location of the bacteria. The edge colors represent positive (green) and negative (orange) correlations. Only significant interactions are shown (r > 0.7). The motile Bacillus in the bulk soil are separated from other bacteria.

### 3.5 Plant trait data

Without the 0.45-µm filter membrane, plant growth differed under different nitrogen concentrations. The *spinach* in H2 was larger than the *spinach* in H1 and H3, and the *spinach* fresh weight in H2 was 23.3% and 14.6% higher than the weights in H1 and H3, respectively. After filter membrane addition, there was no significant difference in *spinach* growth among the three nitrogen treatments. Moreover, *spinach* grown without the filter membrane coating grew better than *spinach* grown with the filter membrane coating. The fresh weights of *spinach* grown without the filter membrane were 48.5% (H1-NF versus H1-F), 56.2% (H2-NF versus H3-F), and 57.8% (H3-NF versus H3-F) greater than those grown with the filter membrane addition (Fig. 6).

**Figure 6.**
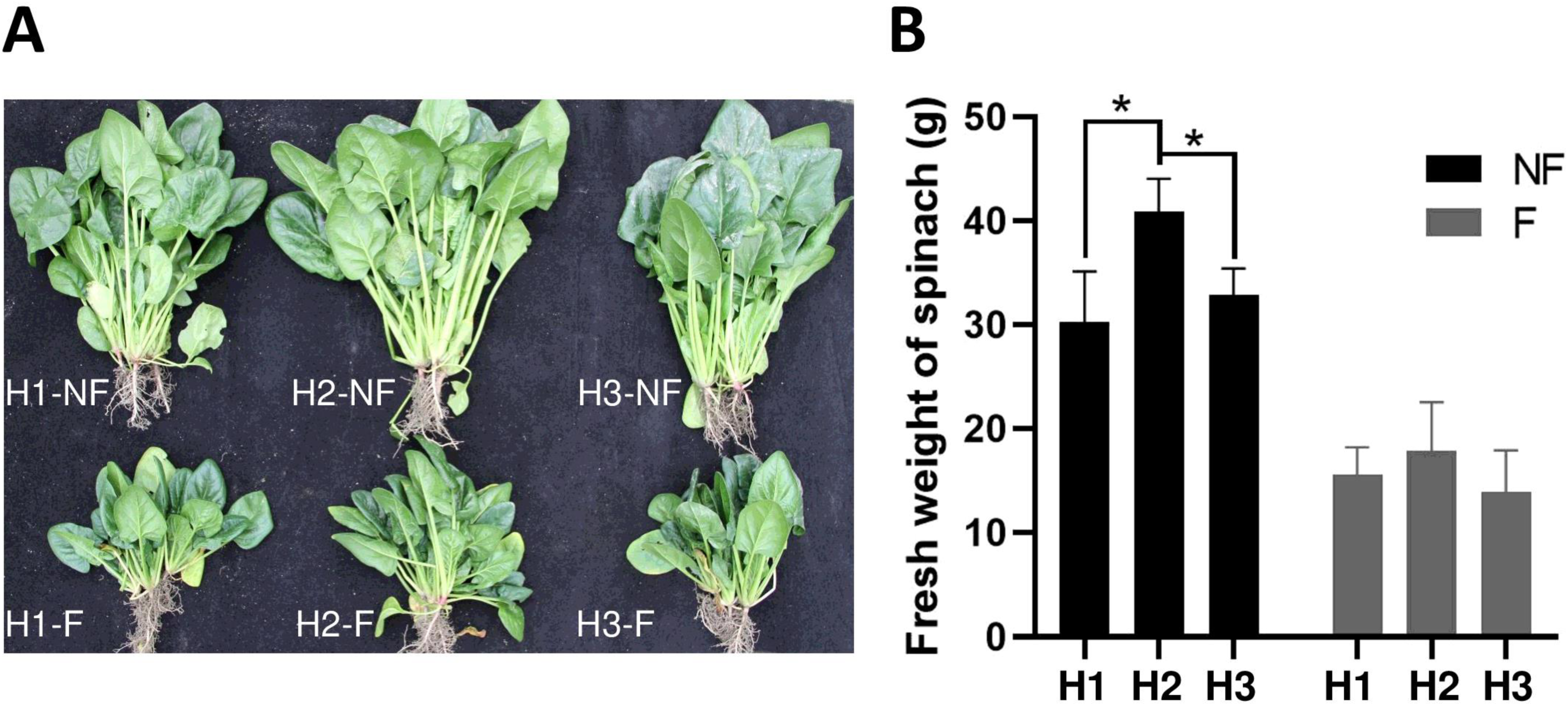
Comparison of spinach growth under different nitrogen concentrations (H1–3), with (F) or without (NF) the filter membrane. (A) Photos of mature spinach; (B) Statistical analysis of spinach fresh weights. Error bars are standard errors, and asterisks indicate significance at **P < 0.01. H1, H2, and H3, normal, normal + 20%, and normal + 40% nitrogen fertilizer concentrations, respectively.

## 4. Discussion

Microorganisms in the soil play significant roles in plant development(Sheng et al., 2021; Sun et al., 2021; Warmink and van Elsas, 2009; Zhao et al., 2014). They are an essential component of soil ecology and serve as an important mechanism for soil regeneration. However, culture-independent approaches have shown that the microbial diversity levels of the soil and rhizosphere are highly underestimated(Chaparro et al., 2014; Korenblum et al., 2020; Wang et al., 2020). Moreover, the driving forces behind changes in microbial diversity are unclear. Our research found that hitchhiking strongly affects bacterial community structure, which is strongly regulated by nitrogen concentration and plants.

### 4.1 Plant and nitrogen concentrations affect microbial composition, especially *Bacillus*

From the analysis of microbial composition, *Bacillus* was found to have no significant changes among the different nitrogen concentrations in the groups without plants, which showed that the content of *Bacillus* could not be affected by nitrogen fertilizer. In groups with plants, when 20% nitrogen fertilizer was added, the content of *Bacillus* was highest compared with the other two groups, which suggested that nitrogen fertilizer can affect the recruitment of *Bacillus*. Under different nitrogen concentrations, the quality of root exudates, such as terpenoid secretion, often changed, which was previously believed to help root colonization of *Bacillus*. Furthermore, this trend was prevented by the filter membrane, which indicated that the filter membrane can prevent the motility of *Bacillus*.

Similar to the contents of *Bacillus*, there was no significant difference in beta diversity when plants were absent. With plants, the bulk soil and rhizosphere microbial community structure of the groups without filter membranes showed significant beta-diversity levels. However, the addition of the filter membrane eliminated the effects of different nitrogen fertilizer treatments on the community structure. These results suggest that nitrogen fertilizer and plants synergistically regulate soil and rhizosphere microbial communities through microbial migration. This is consistent with previous studies on the driving forces of prokaryotic community change, which concluded that soil physical and chemical properties affect cellular dynamics and then microbial communities(Mitchell and Kogure, 2006; Torsvik et al., 2002).

### 4.2 Plant and nitrogen concentrations have important effects on cell motility

If there were no plants in the soil, the functional abundance of cell motility showed no great difference between different nitrogen concentrations. This indicated that nitrogen had no effect on cell motility alone. When plants are present, nitrogen concentration has a greater effect on cell motility. Moreover, the addition of the filter membrane weakened the variation in the functional abundance of cell motility among different nitrogen concentrations, which corresponds to the content of *Bacillus* under different conditions. These results suggest that nitrogen fertilizer and plants synergistically regulate microbial motility. Especially when plants have no membrane, cell motility and *Bacillus* of H2 are the highest compared with other plants. Nitrogen can trigger genes related to motility in motile bacteria and rewiring of the nitrogen regulation system(Karthikeyan et al., 2021; Kavamura et al., 2018; Sun et al., 2020; Wang et al., 2021; Zhou et al., 2017). At the same time, plant secretions were different under different concentrations. Under appropriate nitrogen concentrations, plant secretions might have higher sensitivity to *Bacillus*(Yuan et al., 2015).

### 4.3 The effect of microbial hitchhiking was promoted by a proper concentration of nitrogenous fertilizer

Studies using microscopy methods, motility assays, and genetic approaches have shown that *Bacillus* transports *Streptomyces spores* by direct attachment (hitchhiking) to the bacterial flagella to utilize plant root exudate as a nutrient source(Muok et al., 2021). Based on this, *Bacillus* was selected to explore whether its interactions with other bacteria were stronger. We discovered that more bacteria were impacted by the motile *Bacillus* in H1, and this effect was also observed in H3 but not in H2. In the membrane-addition treatment group, there were fewer interactions between motile bacteria in the rhizosphere and soil under the three different nitrogen concentrations. Thus, the presence of a filter membrane precluded the interactions between soil and rhizosphere microorganisms mediated by motile bacteria. We concluded that changes in nitrogen concentration affect the motility of and hitchhiking onto motile bacteria.

The co-occurrence network, which provides insights into ecological interactions among microbial taxa, shows that the interactions between different microbial strains are the main driving factors of population structure and dynamics because microbes can co-occur or exclude each other, revealing that different microorganism metabolites can have an inhibitory or symbiotic impact. Co-occurrence network analyses have shown that taxonomic shifts owing to the variation in nitrogen availability can shape patterns of ecological interactions regulating the structure, function, and potential resilience of soil microbial communities (Han et al., 2020; Schmidt et al., 2019). However, these networks have focused only on the soil or rhizosphere microbiome and did not take into account the interactions between these two communities. Our cooccurrence network analysis combining soil and rhizosphere microbes showed that there were variations in the cooccurrence of bulk soil and rhizosphere microbial species at varying nitrogen concentrations.

### 4.4 Nitrogen influenced plant growth by hitchhiking

Morphological variations in plants may result from different chemical property(Busoms et al., 2021) especially fertilizer concentrations in soil (Sun et al., 2021). Fertilization with an appropriate nitrogen content promotes plant growth, while fertilization with a high nitrogen content inhibits plant growth, which is consistent with our results (Bindraban et al., 2020; Bright et al., 2021; Elhanafi et al., 2019; Liu et al., 2014; Razaq et al., 2017). Previous studies have suggested that root length and root surface area increased under intermediate N levels and that root growth was reduced under both higher and lower fertilization levels, affecting plant morphology(Chen et al., 2020; Good and Beatty, 2011). On the other hand, nitrogen is functional in the construction of and chlorophyll, which can influence plant growth and development by affecting photosynthesis and the uptake of minerals(Grossart et al., 2001; Kavamura et al., 2018; Razaq et al., 2017; Wang et al., 2021). In addition to these reasons, in our study, we found that in the treatment group with filter-membrane addition, there was no significant difference in the growth of *spinach* with different nitrogen concentrations. Through correlation analysis, we found that the hitchhiking phenomenon was blocked by the filter membrane; as a result, we presumed that nitrogen may affect plant growth by affecting hitchhiking. In the rhizosphere, compared with H1 and H3, there was an increase in functional abundance encouraging plant growth (amino acid metabolism, energy metabolism, and cofactor and vitamin metabolism) and a decline in functional abundance (infectious disease) discouraging plant growth in the H2 group without a filter membrane. At suitable concentrations, the nonmotile bacteria brought by the hitchhiking phenomena modified the rhizosphere microbial environment via transport on favorable bacterial species, resulting in improved plant growth. It can be concluded that nitrogen fertilizer can promote plant growth by promoting microbial hitchhiking.

## 5. Conclusion

Nitrogen alone has no effect on hitchhiking, but it will synergistically affect the movement of sports bacteria with plants, thus causing the carrying of sports bacteria. Moreover, different concentrations of nitrogen fertilizer have different effects on hitchhiking. The appropriate concentration will greatly improve, while an excessive concentration will reduce its effect. These studies provide guidance for rational fertilization.

## Acknowledgments

This project was funded by the National Natural Science Foundation of China (NSFC Grant No. 31870240 and No. 51909175), the Key Research and Development Program of Sichuan Province (No. 2019YFN0153) and the Modern Agriculture Discipline Construction and Promotion Program of Sichuan Academy of Agricultural Sciences (2021XKJS032). We thank Mr. Mingyuan Wang for his guidance and advice during the preparation of this manuscript.

## Appendix A. Supplementary material

Supplementary material.

## Author contributions

LZB, YL, LC, YY and TX designed the experiment; CW, ZQ, KWQ, WXB, and LZ conducted the experiments; WZY and XQN analyzed the data; WZY, LWY, LXR and WYF wrote the manuscript.

## Ethics approval

Not applicable.

## Conflict of interest

The authors declare no competing interests.

---

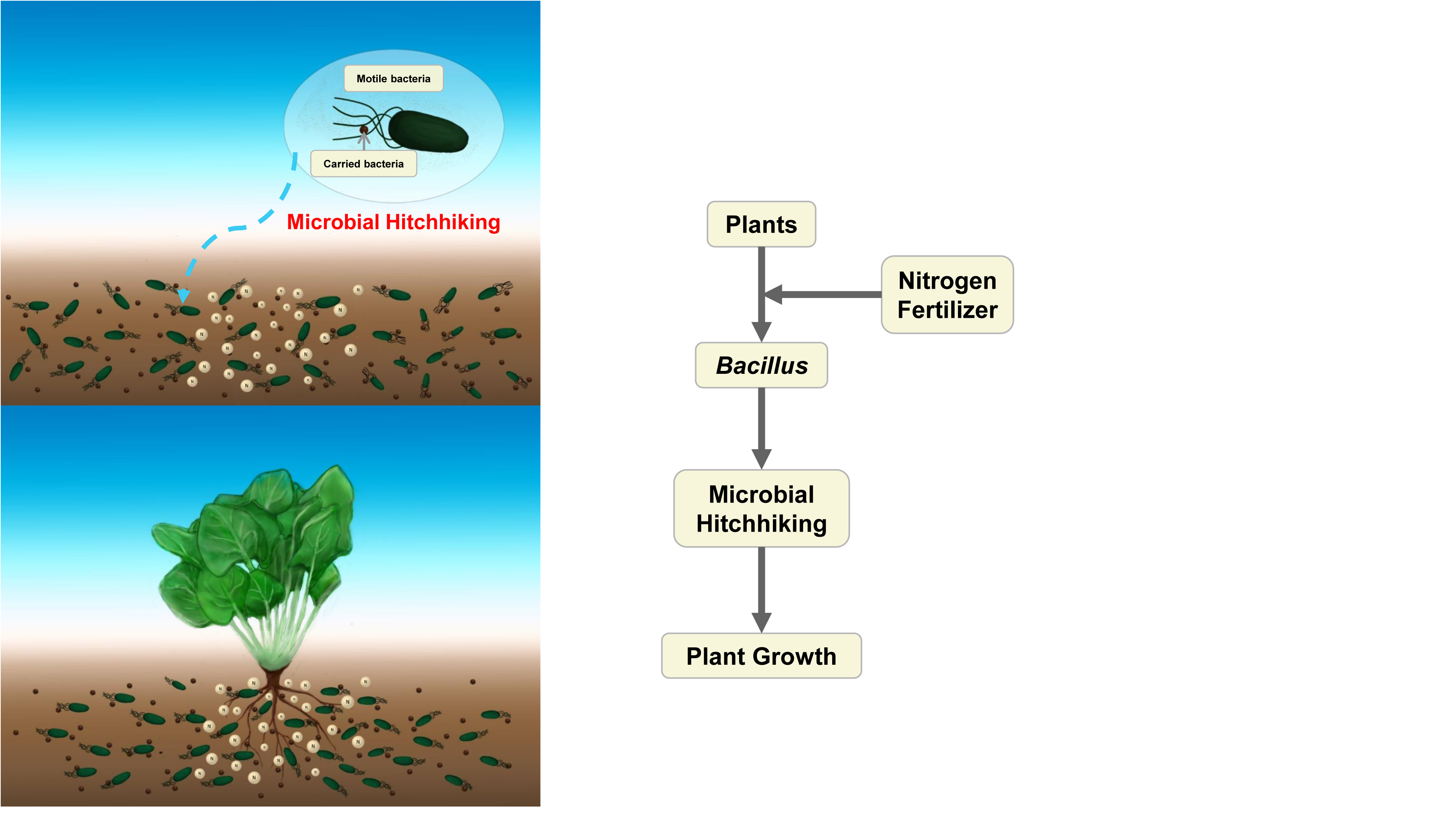

## Notes

### Competing Interest Statement

The authors have declared no competing interest.

