## Supplementary material for "The synergistic effects of plants and nitrogen on microbial hitchhiking": supp: Supplementary Figures.pptx

### Slide 1
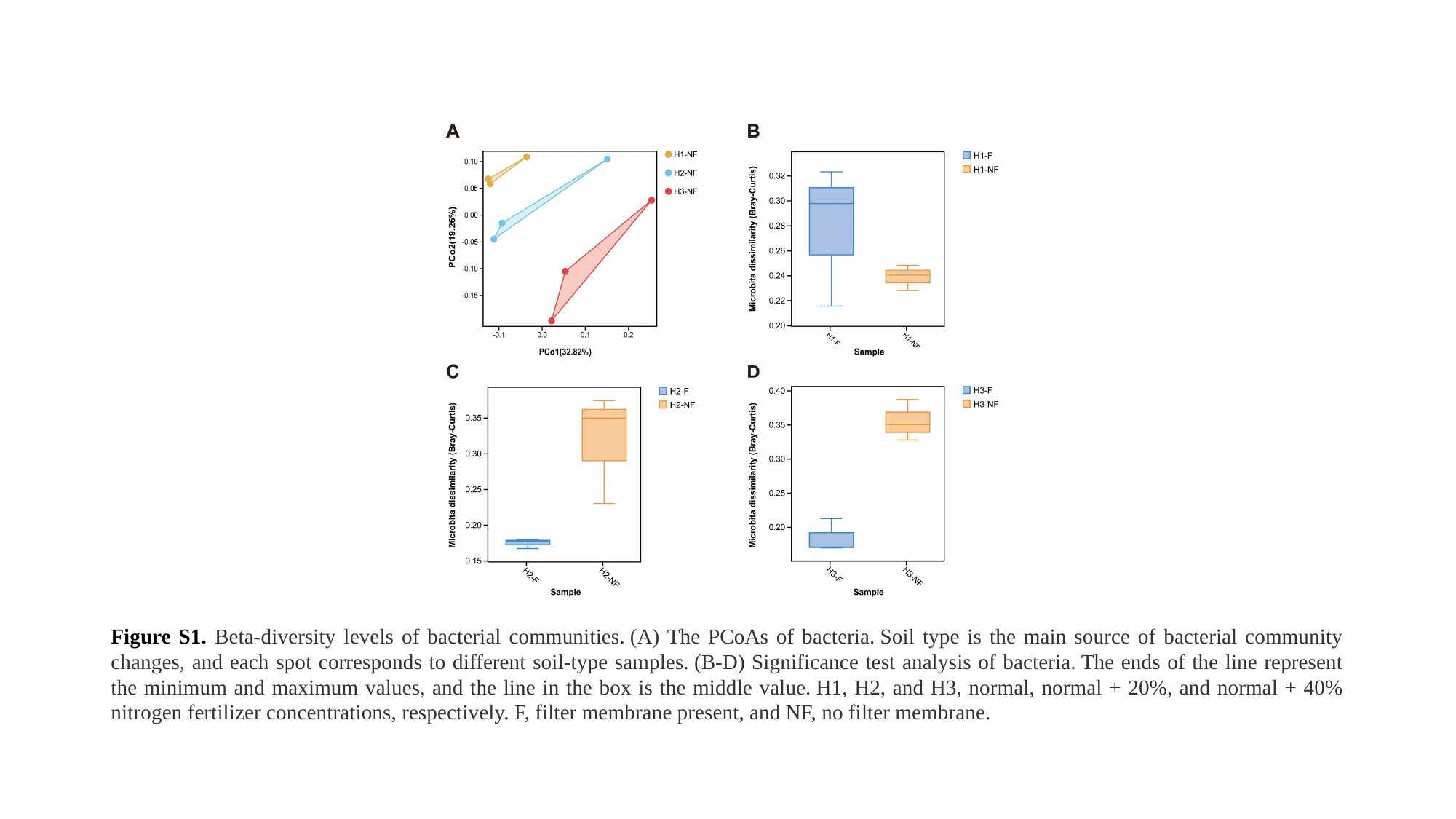

#### Figure S1. Beta-diversity levels of bacterial communities. (A) The PCoAs of bacteria. Soil type is the main source of bacterial community changes, and each spot corresponds to different soil-type samples. (B-D) Significance test analysis of bacteria. The ends of the line represent the minimum and maximum values, and the line in the box is the middle value. H1, H2, and H3, normal, normal + 20%, and normal + 40% nitrogen fertilizer concentrations, respectively. F, filter membrane present, and NF, no filter membrane.

### Slide 2
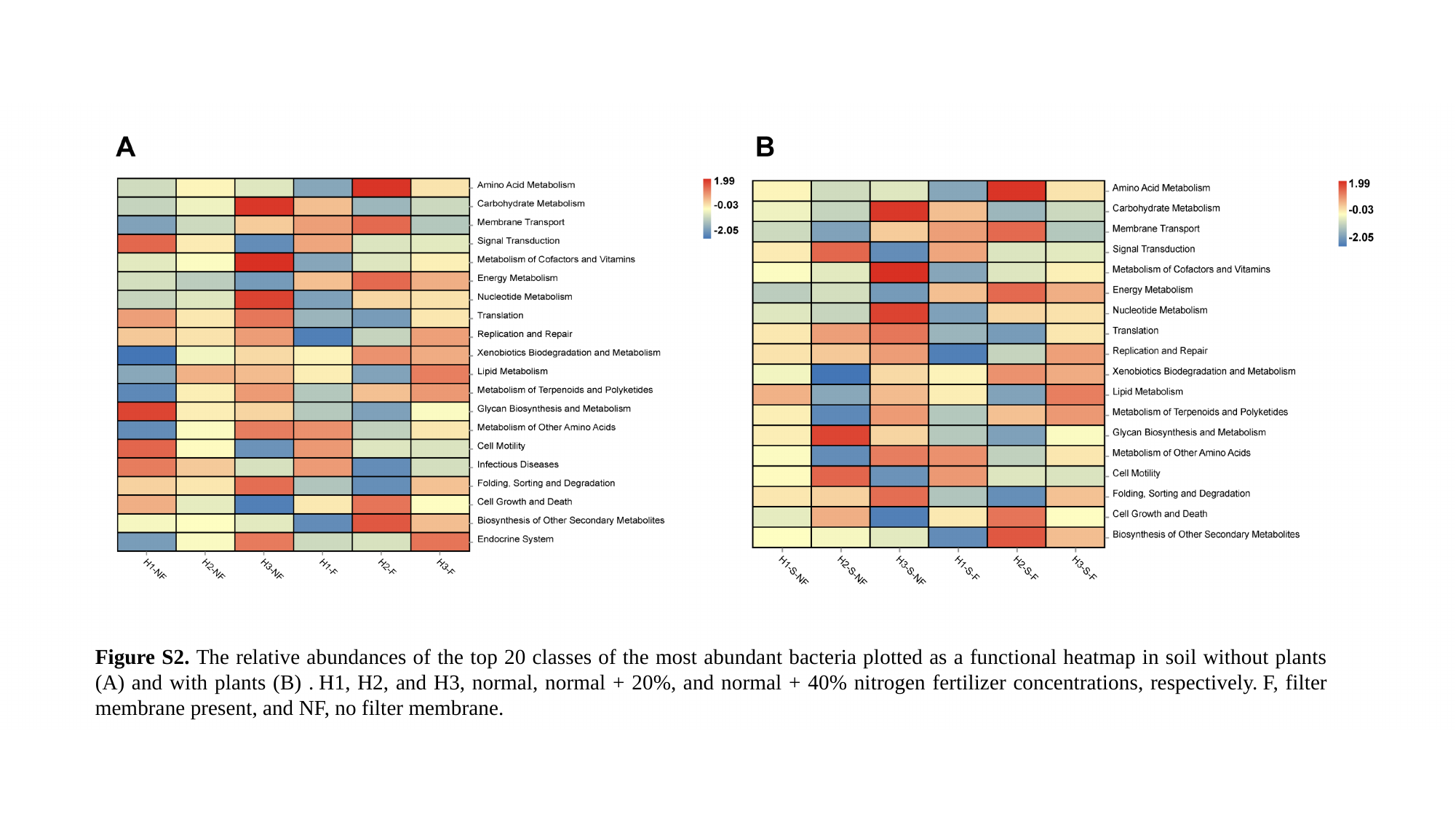

#
Figure S2. The relative abundances of the top 20 classes of the most abundant bacteria plotted as a functional heatmap in soil without plants (A) and with plants (B) . H1, H2, and H3, normal, normal + 20%, and normal + 40% nitrogen fertilizer concentrations, respectively. F, filter membrane present, and NF, no filter membrane.
