## Supplementary material for "The synergistic effects of plants and nitrogen on microbial hitchhiking": supp: Supplementary Information.docx

RESEARCH PAPER

**Soil nitrogen impacts the rhizosphere microbial community owing to microbial hitchhiking**

^5^ Guangdong Provincial Key Laboratory of Silviculture, Protection and Utilization, Guangdong Academy of Forestry, 233 Guangshan 1st Road, Guangzhou, PR China.

**Contents**

**Supplementary Materials and Methods**

DNA extraction and PCR amplification

Illumina Novaseq 6000 sequencing

Quality control and clustering

**Supplementary Tables**

**Table S1** Abundances of class-level microorganisms in all groups;

**Table S2** Abundances of OTUs in all groups;

**Table S3** Alpha-diversity in all groups (sobs, shannon, simpson, chao, ace, goods, coverage, pielou and pd index);

**Table S4** Determinations of the alpha- and beta-diversity levels in the bacterial and fungal communities under different nitrogen concentrations. A) Kruskal–Wallis H test results for bacterial alpha-diversity levels; B) Anosim results for bacterial beta-diversity levels;

**Table S5** Functional abundance for all groups;

**Table S6** Primers used in this study for quantitative real-time polymerase chain reaction (qPCR).

**Supplementary Figures**

**Figure S1** Beta-diversity levels of bacterial communities. (A) The PCoAs of bacteria. Soil type is the main source of bacterial community changes, and each spot corresponds to different soil-type samples. (B-D) Significance test analysis of bacteria. The ends of the line represent the minimum and maximum values, and the line in the box is the middle value. H1, H2, and H3, normal, normal + 20%, and normal + 40% nitrogen fertilizer concentrations, respectively. F, filter membrane present, and NF, no filter membrane.

**Figure S2** The relative abundances of the top 20 classes of the most abundant bacteria plotted as a functional heatmap in soil without plants (A) and with plants (B) . H1, H2, and H3, normal, normal + 20%, and normal + 40% nitrogen fertilizer concentrations, respectively. F, filter membrane present, and NF, no filter membrane.

**Supplementary Materials and Methods**

*The map of pots in the greenhouse*

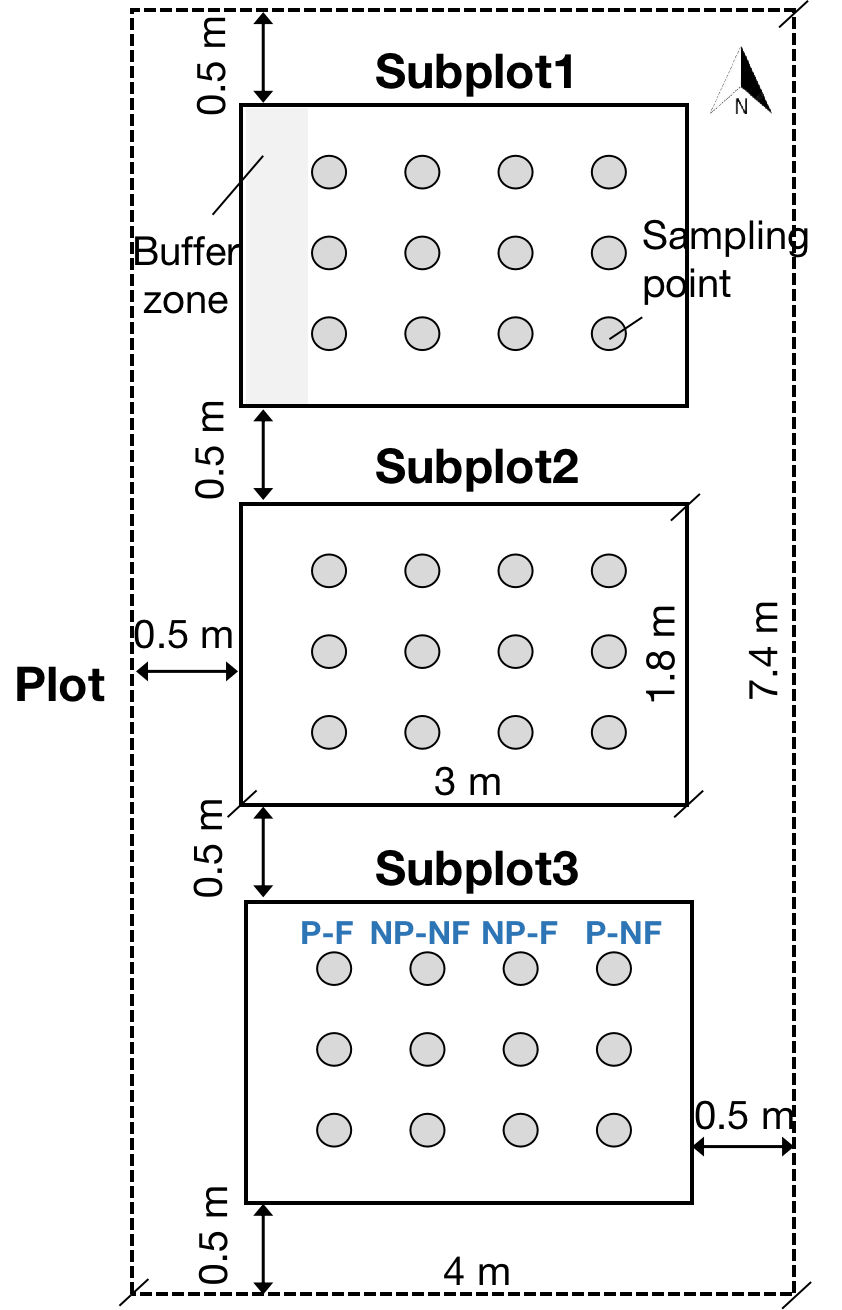

Assignment of pots in the greenhouse. P-F denotes treatment with plant in coated pot; NP-NF denotes treatment without plant in uncoated pot

*DNA extraction and PCR amplification*

Microbial DNA was extracted using the HiPure Soil DNA Kits (or HiPure Stool DNA Kits ) (Magen, Guangzhou, China) according to manufacturer’s protocols. The 16S rDNA target listed in the table

region of the ribosomal RNA gene was amplified by PCR 94 °C for 2 min, followed by 30 cycles at 98 °C for 10 s, 62 °C for 30 s (except for 16S V4: 55°C for 30 s), and 68 °C for 30 s and a final extension at 68 °C for 5 min using primers listed in the table (Guo et al., 2017). PCR reactions were performed in triplicate 50 μL mixture containing 5 μL of 10 × KOD Buffer, 5 μL of 2 mM dNTPs, 3 μL of 25 mM MgSO4, 1.5 μL of each primer (10 μM), 1 μL of KOD Polymerase, and 100 ng of template DNA. Related PCR reagents were from TOYOBO Japan. Other primers are listed in Table S6.

*Illumina Novaseq 6000 sequencing*

Amplicons were extracted from 2% agarose gels and purified using the AxyPrep DNA Gel Extraction Kit (Axygen Biosciences, Union City, CA, U.S.) according to the manufacturer’s instructions and quantified using ABI StepOnePlus Real-Time PCR System (Life Technologies Foster City, USA). Purified amplicons were pooled in equimolar and paired-end sequenced (PE250) on an Illumina platform according to the standard protocols.

*Quality control and clustering*

***Reads filtering***

Raw data containing adapters or low quality reads would affect the following assembly and analysis. Thus, to get high quality clean reads, raw reads were further filtered according to the following rules using FASTP(Chen et al., 2018) (version 0.18.0):

1) Removing reads containing more than 10% of unknown nucleotides (N);

2) Removing reads containing less than 50% of bases with quality (Q- 20.

***Reads assembly***

Paired end clean reads were merged as raw tags using FLSAH (Magoč and Salzberg, 2011) (version 1.2.11) with a minimum overlap of 10 bp and mismatch error rates of 2%.

***Raw tag filtering***

Noisy sequences of raw tags were filtered under specific filtering conditions to obtain the high-quality clean tags. The filtering conditions are as follows

1) Break raw tags from the first low quality base site where the number of bases in the continuous low quality value (the default quality threshold is 3) reaches the set length (the default length is 3

2) Then, filter tags whose continuous high-quality base length is less than 75% of the tag length.

2.1.4 Clustering and chimera removal

The clean tags were clustered into operational taxonomic units (OTUs) of ≥≥ 97 % similarity using UPARSE(Edgar, 2013) (version 9.2.64) pipeline. All chimeric tags were removed using UCHIME algorithm (Edgar et al., 2011) and finally obtained effective tags for further analysis. The tag sequence with highest abundance was selected as representative sequence within each cluster.
